# Decoding Fear or Safety and Approach or Avoidance by Brain-Wide Network Dynamics

**DOI:** 10.1101/2022.10.13.511989

**Authors:** Danilo Benette Marques, Matheus Teixeira Rossignoli, Bruno de Avó Mesquita, Tamiris Prizon, Leonardo Rakauskas Zacharias, Rafael Naime Ruggiero, João Pereira Leite

## Abstract

Discerning safety from threat and positive or negative outcomes of adversity are fundamental for mental health. Many brain structures have been implicated in both adaptive and maladaptive stress coping, however, how multiple regions function together as a network in the processing of this information is unclear. Here, we recorded local field potentials from seven regions of the mesolimbic-hippocampal-prefrontal cortical network (MLHFC) of male rats during the conditioning of a stimulus (CS) to the absence (safety) and then to the anticipation (fear) of footshocks, and during an approach-avoidance task. We developed a machine learning pipeline to investigate the relevance of specific features of oscillatory activity in the decoding of fear versus safety and approach versus avoidance. We found that decoding performance increased as a function of the number of brain regions included, reaching the best classification if all regions were considered. In addition, the best decoding was obtained from frequencies within the theta range (4-10 Hz). Remarkably, decoder models showed robust generalization within but not between individuals. Nevertheless, we were also able to identify patterns of MLHFC activity that decoded stress coping states from all rats. These patterns were characterized by increased brain-wide theta synchrony during fear and preceding approach. Our results indicate that stress coping information is encoded at the brain-wide level and highlight individual variability in this neural processing. Our findings also suggest that MLHFC network theta activity underlies active stress coping with both aversive and positive motivational valences.

**SIGNIFICANCE STATEMENT:** The appraisal of safety versus threat and positive versus negative valence of adversity are core dimensions of emotional experience and stress coping. We developed a new behavioral protocol that discriminates states of fear, safety, approach, and avoidance in a single subject and a machine learning-based method to investigate how neural oscillations can decode such states. Our work provides evidence that stress coping is processed at multiple regions on a brain-wide level involving network oscillations at the theta frequencies, which especially synchronizes during fear and approach. We highlight the potentials of combining artificial intelligence and multi-site electroencephalography to guide therapeutic decisions in precision psychiatry and theta-boosting stimulation therapies for stress-related disorders, especially related to cognitive and motivational deficits.

## Introduction

The ability to differentiate between safe and stressful situations and the accurate appraisal of positive or negative outcomes from adversities are fundamental for survival and mental health (Kong et al., 2014; Southwick et al., 2015). Imbalances of these appraisals towards threat and aversion are hallmarks of psychiatric illnesses, such as post-traumatic stress disorder, generalized anxiety disorders, and major depression (Trew, 2011; Jovanovic et al., 2012; Gazendam et al., 2013). In turn, fostering stress regulation and positive emotions are therapeutic goals for these disorders and improve overall quality of life (Feder et al., 2019).

Many brain regions, including the prefrontal cortex, the hippocampus, the amygdala, and the mesolimbic reward system, have been implicated in processing adaptive and maladaptive stress coping (Godoy et al., 2018; Ruggiero et al., 2021). Neural oscillations have been suggested to bind the activity of distant areas and underlie the encoding of information (Fries, 2015; Harris and Gordon, 2015), where the theta oscillations (4-12 Hz) are the most widely associated with cognition and emotion (Korotkova et al., 2018). Previous studies reported increases in limbic-cortical theta power and synchrony during avoidance in the elevated plus maze (Adhikari et al., 2010; Jacinto et al., 2016) and conditioned fear (Seidenbecher et al., 2003; Taub et al., 2018), implicating this activity in the “aversive” component of stress (Çalışkan and Stork, 2019). However, recent evidence indicates that hippocampal-prefrontal theta activity during fear is related to the expected controllability over stressors (Marques et al., 2022), which is associated with resilience (Maier et al., 2006). In addition, changes in prefrontal-amygdala theta directionality are associated with safety and fear extinction (Lesting et al., 2013; Likhtik et al., 2014). Furthermore, past investigations did not make explicit the rewarding component of stressful experiences. Therefore, how these activities are related to positive valence is unknown. A clearer dissection of safety and valence would elucidate the participation of theta activity in stress coping.

Although specific regions and projections have been implicated, it is theorized that multiple brain structures function together as a network in processing stress responses (Sousa, 2016; Grossman and Dzirasa, 2020). However, how these regions interact is unclear. Machine learning is an emergent and promising approach for identifying complex multivariate patterns of neural network activity and individual differences in brain function related to depression and resilience (Drysdale et al., 2017; Dunlop et al., 2017; Vu et al., 2018). Recently, using machine learning, Hultman et al. (2018) successfully identified a brain-wide oscillatory pattern predicting resilient versus susceptible mice to social stress. However, the significance of such patterns remains unclear, and the replicability of this approach is still a matter of debate.

To address these issues, we recorded local field potentials from seven regions of the mesolimbic-hippocampal-prefrontal cortical network (MLHFC) of rats in a new experimental design where an identical stimulus is conditioned to safety and later to fear and these associations are tested in an approach-avoidance task with explicit reward, aversive and safe locations. Then, using machine learning, we investigated the performances of distinct features of oscillatory activity in decoding fear or safety and approach or avoidance. We hypothesized that neural decoding of stress states would perform better if the oscillatory activity of all regions recorded were used together. We observed that decoding performance improved as a function of the number of regions included in the model, achieving the highest performance if all regions were included. We also found that, across all frequencies, the best decoding happened within the theta range. Noteworthy, the decoder models exhibited generalization within each rat but not between individuals, demonstrating relevant individual variability in stress processing. Nevertheless, our algorithm discovered patterns of brain-wide theta synchrony that decoded periods of fear and preceding approach across all rats. Our results demonstrate that stress coping is encoded at the brain-wide level and suggest that MLHFC network theta activity underlies active stress coping with both aversive and, especially, positive motivational valence.

## Materials and Methods

### Subjects

Adult male Sprague-Dawley rats (8-10 weeks old weighing 320-360 g, Ribeirão Preto, Brazil) were housed singly in bedded cages in a temperature controlled room (22 ± 2 °C) on a 12 h light/dark cycle (lights on at 7 A.M.) with *ad libitum* access to food and water. The procedures followed the National Council for the Control of Animal Experimentation guidelines and were approved by the local Committee on Ethics in the Use of Animals (Ribeirão Preto Medical School, University of São Paulo, protocol 0004/2020).

### Electrode implantation surgery

Rats were anesthetized with ketamine and xylazine (respectively: 50 mg/Kg and 25 mg/Kg intraperitoneal, followed by 70 mg/Kg and 35 mg/Kg intramuscular). Body temperature was kept constant during the entire procedure by a heating pad (37 ± 1 °C).

Multi-site electrode systems consisted of an eight-channel connector (Omnetics), with each channel soldered to a bare silver wire (A-M Systems, 381 µm). Monopolar electrodes (teflon-coated tungsten, 50 µm) were lowered into each region, fixed with dental cement and soldered to a silver wire. The targeted bregma-referenced coordinates were: prefrontal cortex prelimbic (PL; anterior: 3 mm, lateral: 0.5 mm, ventral: 3.2 mm) and infralimbic areas (IL; anterior: 3 mm, lateral: 0.5 mm, ventral: 4.2 mm), nucleus accumbens shell (NAc; anterior: 1.7 mm, lateral: 0.6 mm, ventral: 6.8 mm), basolateral amygdala (AMG; posterior: 2.6 mm, lateral: 5.1 mm, ventral: 6.7 mm), dorsal hippocampus CA1 (DH; posterior: 4.2 mm, lateral: 4 mm, ventral: 2.8 mm), ventral hippocampus CA1 (VH; posterior: 4.8 mm, lateral: 5 mm, ventral: 7.5 mm), and ventral tegmental area (VTA; posterior: 5.8 mm, lateral: 0.8 mm, ventral: 8.2 mm) (Paxinos and Watson, 2009; Bueno-Junior et al., 2018; Ruggiero et al., 2018; Marques et al., 2022).

In addition to the recording electrodes, microscrews were fastened into the skull, including a ground reference in the right occipital bone close to the lambda. Electrodes and screws were then cemented together with acrylic resin. Analgesic, antipyretic, and antibiotic drugs were injected after surgery. Animals were allowed to recover for 8-9 days before food restriction.

### Behavioral apparatus

#### Shuttle box

We used a customized shuttle box system for simultaneous electrophysiological recording around shocks and behavioral monitoring (Marques et al., 2022). The behavioral apparatus was located inside a soundproof box and consisted of a chamber (54 cm length × 33 cm width × 50 cm height) divided into two compartments by a removable wall (1 cm length × 1.5 cm height). Footshocks were delivered through stainless steel bars on the floor. The apparatus contained eight equally spaced infrared beams, four per compartment, to track position and record crossings.

#### Reward and platform devices

We customized a reward delivering device and a footshock-avoiding platform with presence sensors. The reward device was developed over a bird feeder with a single opening (3 cm length × 1.9 cm width × 1.7 cm height) that allowed the entry of the rostral part of the rat’s head. The feeder was attached to a 50 ml falcon tube filled with 50 ml of a 20% sucrose solution before each session. We soaked the device on vanilla odor to serve as a cue. Two infrared emitter-sensor 0.3 mm LED pairs were fixed in the laterals of the feeder to detect the rat’s head during reward consumption. Before each approach-avoidance test, the reward device was fixed on the left wall of the shuttle box, while the platform (8 cm length × 16 cm width × 1.5 cm height) was placed in the right corner. A single infrared emitter-sensor 0.5 mm LED pair was fixed outside the shuttle box with the beam aimed at the center of the platform to detect the rat’s presence. Blocking of infrared light beams was detected with Arduino Uno (Arduino), which sent timestamps at 1 kHz to the electrophysiological recording system.

### Experimental design

All behavioral procedures were performed during the light phase (8 A.M. to 6 P.M.) in a temperature-controlled (22 ± 2°C) dark room (0 lux). After surgery recovery, animals underwent food restriction (∼20 g of chow biscuits per day) to 90% of body weight. Animals were habituated to the behavioral apparatus, the reward, and the platform for 30 min on the first day and to the conditioned stimulus on the following day (Fig. 1*A*). All animals consumed the reward during habituation.

**Figure 1.**
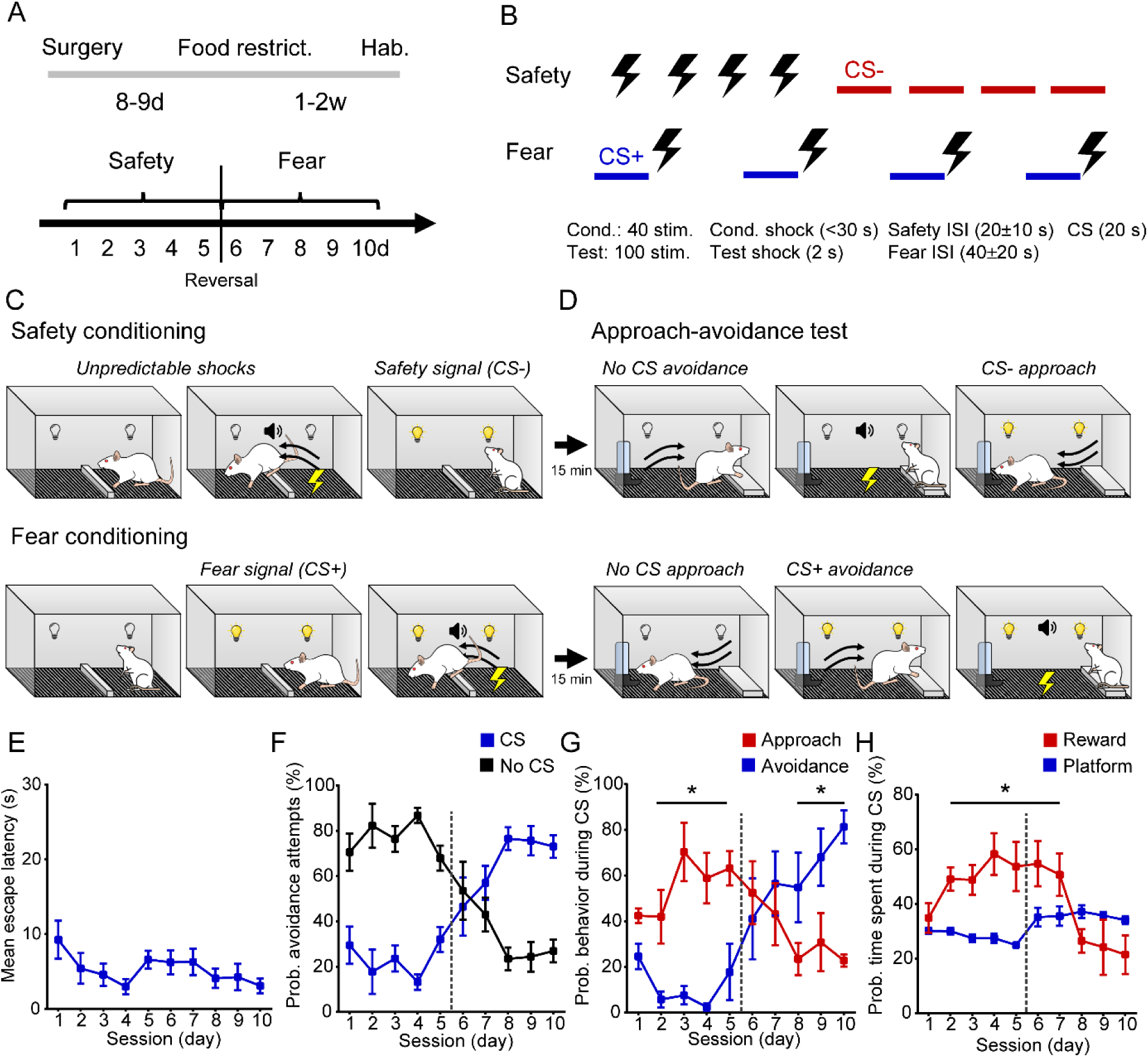
Differential conditioning of the same stimulus to safety or fear promotes approach or avoidance in a within-subject design. ***A*,** Experimental design. Chronically implanted animals are food-restricted and submitted to days of safety and then fear conditioning and testing. ***B*,** Contingencies between conditioned stimulus (CS) and shocks for safety and fear conditioning. In the safety protocol, shocks and CS were presented separately, while in the fear protocol the CS always preceded escapable shocks. ***C-D*,** Graphical representation of behavioral results in safety (upper) and fear (lower) protocols of conditioning **(*C*)** and test **(*D*)**. ***E*,** Constant average escape latencies to shocks throughout conditioning sessions. ***F*,** CS-suppresses avoidance attempts, and CS+ increases them in conditioning sessions. ***G-H*,** Safety conditioning promotes higher probabilities of approach and time on reward, while reversal to fear gradually increases the probabilities of avoidance and time on platform in the approach-avoidance test. CS = conditioned stimulus; ISI = interstimulus interval. *p < 0.05 (Fisher’s LSD test).

The present experimental design is a merge of the learned safety (*versus* learned fear) (Pollak et al., 2010a) and the platform-mediated avoidance (Bravo-Rivera et al., 2014; Diehl et al., 2019) protocols, adapted for a within-subject investigation (Fig. 1*A-D*). Each day consisted of a conditioning session (Fig. 1*C*), followed by an approach-avoidance (AA) test (Fig. 1*D*). On the initial days, we conditioned a signal to the absence (safety) of aversive stimuli and, on the following days, to the anticipation (fear) of them. Then, to reveal these associations, we expected the safety signal to increase approach to the reward, while the fear signal would increase avoidance to the safe platform in the AA test (Fig. 1*C-D*).

The conditioned stimulus (CS) was a LED light (200 lux) of 20 s. The aversive unconditioned stimulus (US) was an oscillating footshock of 0.6 mA during conditioning sessions and 0.8 mA in the AA test. All shocks were concomitant to a sound stimulus (60 dB tone). During conditioning, shocks were escapable and would terminate when the rat crossed between compartments or after 30 s. During AA test, shocks had a fixed duration of 2 s and could be avoided by standing on the platform.

On the first 3-5 days, the CS was conditioned and tested for safety (CS-) by explicitly unpairing it with the US. In the safety protocol, the interstimulus interval (ISI) was random in the range of 20 ± 10 s. The US and CS were presented 4 times each for 10 times (40 trials) during conditioning and 25 times (100 trials) in the AA test (Fig. 1*B*). After the last safety session, there was a reversal of the CS association. On the 3-5 following days, the CS was conditioned and tested for fear (CS+). In the fear protocol, the CS preceded and accompanied every US, followed by a 40 ± 20 s ISI. The CS-US pairings were presented 40 times during conditioning and 100 times in the AA test (Fig. 1*B*). After each conditioning session, the wall was removed, and the reward and platform were inserted. The reward compartment was blocked until the start of the AA test after 15 min (Fig. 1*C-D*).

We quantified the shock escape latencies and the number of crossings during and out of CS (avoidance attempts) in the conditioning sessions. In the AA test, we recorded the crossings to the reward compartment (approach) and to the platform compartment (avoidance). Then, we quantified the probability of these behavioral responses during CS (number of responses during CS/total number). We also measured the time spent in the reward and on the platform and the probability of time spent in these locations during CS (time spent in local during CS/total time spent in local).

### Extracellular recordings

Electrophysiological signals were recorded during conditioning and test sessions, interrupted only during footshocks by a relay system. A multichannel acquisition processor (Plexon) was used to record local field potentials (LFP) with the following parameters: 1000x gain, 0.7-500 Hz band-pass filter, 1 kHz sampling rate. Timestamps were acquired from the behavioral apparatus at 40 kHz sampling.

### Histology

After the last session, animals were euthanized with CO2 asphyxiation, and electrolytic lesion currents (1 mA) were applied between pairs of wires. After decapitation, each brain was immersed in 4% paraformaldehyde overnight (PFA, −20 °C), followed by 70% ethanol and paraffin for coronal sectioning at the microtome. Standard cresyl-violet staining was used to validate electrode positioning under the bright-field microscope.

### Data analysis

Signal processing and analysis of electrophysiological data were performed using custom scripts in MATLAB.

#### Local field potentials

LFPs were epoched in 1-s segments with no overlap (Fig. 2*C*). Epochs with LFP noise (above 0.5 mV) in any region (except the DH) were detected for exclusion. Then, the whole signals were band-pass filtered (1-250 Hz) and concatenated (Fig. 2*D*). For CS analysis, we analyzed the epochs between 1 s and 19 s following CS onset to avoid event-related potentials and shock onset noise. For approach-avoidance (AA) analysis, we analyzed the epochs between 10 s and 1 s before crossings (approach or avoidance). For AA analysis around crossings, we also included 1 s after events. We only considered epochs when the rat was in the opposite compartment prior to each approach (platform side) or avoidance (reward side).

**Figure 2.**
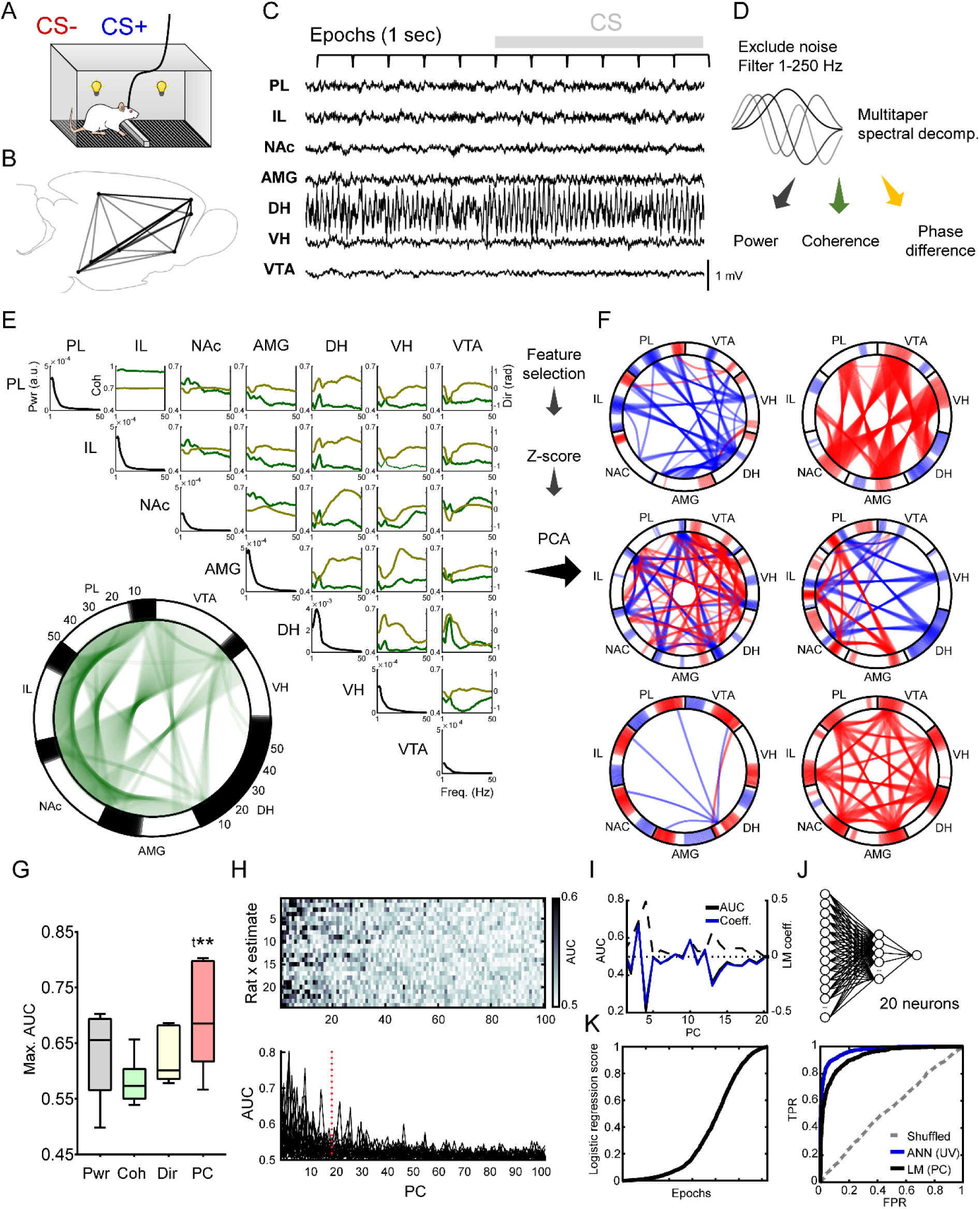
Decoding pipeline. ***A*,** Graphical representation of freely moving electrophysiological recordings during CS- and CS+. ***B*,** Sagittal representation of brain-wide distributed implant coordinates. ***C*,** Representative traces of multi-site local field potentials during CS+. ***D*,** Preprocessing steps and multitaper estimation of spectral power, coherence, and phase difference/directionality. ***E*,** Example of CS+ average raw power (diagonal, black line), coherence (green line), and directionality (yellow line; phase difference) across all regions and pairs of one rat. Network chord diagram (bottom left) showing power (black lines in outer contours) and coherence (green lines within circle) estimates (above a threshold) across frequencies (counterclockwise ascending). ***F*,** After feature selection, the multidimensional spectral estimates are Z-scored and dimensionality-reduced through PCA. Examples of principal components (PC) of different rats. PC coefficients (red = positive; blue = negative) demonstrate brain-wide network collective variations. ***G*,** The average maximum area under the curve (AUC) for CS+/CS-classification by PC scores (from power, coherence, and directionality combined) is greater than by each univariate estimate. ^t^p = 0.058, *p < 0.05 (Fisher’s LSD test). ***H*,** Greater values of AUC occur until the 20th PC for all rats and estimates. ***I*,** Generalized linear model (LM) coefficients correspond to PCs’ AUCs. ***J*,** An artificial neural network with one hidden layer of 20 neurons was also used. ***K*,** Example of LM scores of one rat and its ROC curve (TPR = true positive rate; FPR = false positive rate) compared to shuffled test labels and ANN. Here and on: PL = prelimbic area; IL = infralimbic area; NAc = nucleus accumbens; AMG = amygdala; DH = dorsal hippocampus; VH = ventral hippocampus; VTA = ventral tegmental area.

For each epoch, we used the multi-taper method with 5 tapers and time-half bandwidth product of 3 to obtain spectral estimates of regional power and cross-regional mean phase coherence and mean phase difference. We modified scripts from the Chronux toolbox (Mitra and Bokil, 2009) to obtain coherence estimates equivalent to the phase-locking index described in Lachaux et al. (1999) rather than the magnitude squared coherence because we intended to consider only the time relationships between signals. The phase difference was interpreted as the estimate of directionality between signals (Fig. 2*D*). We calculated spectral data between 1 and 50 Hz using 1024 points for the Fourier transform (50 bins).

### Multivariate analysis and decoding models

#### Decoding pipeline

We developed a decoding pipeline to evaluate how different features of oscillatory activity could classify CS+ *versus* CS-(fear versus safety) or preceding approach (AP) *versus* avoidance (AV) (Fig. 2). The features of interest were: estimate (power, coherence, and directionality); region(s); frequency bin(s); univariate value(s) or multivariate score(s). After selecting the features, each variable was Z-scored throughout epochs of interest, and binary decoder models were generated.

We estimated the binary classification performances of the models’ scores by the area under the receiver operating characteristics curve (AUC-ROC) for each rat. We compared models between features, types, to *chance* (model trained on correct labels but tested on shuffled labels) or to *shuffled* (model trained on shuffled labels and tested on correct labels). Chance and shuffled performances were estimated by the average of 10 randomly shuffling labels.

#### Linear model

The main decoding pipeline used here was a linear model (LM) implemented with the following steps: 1) Feature(s)/attribute(s) selection; 2) Z-score transform based on train data mean and standard deviation; 3) Dimensionality reduction through principal component analysis (PCA) using singular-value decomposition algorithm maintaining the first 20 principal components (PC) unless stated otherwise; 4) Fit of a generalized linear model (GLM) with a binomial distribution and a logit link function (logistic regression) on PC scores (Fig. 2*E*, *F, I, K*).

Except when investigating the relevance of specific datatypes (region, frequency, and estimate), we trained decoder models on all regions, frequencies, and estimates in each rat. For some analyses, when the number of attributes was less than 20 or when we intended to generate an overfitted model, we fitted the GLM on the univariate Z-scored data (UV; LMUV) rather than on PC scores (LMPC).

#### Decoding by region and estimate

To evaluate the performance of a single region, we used the LM on power (of that region), coherence, or directionality (between that region and every other; 6 pairs). To examine how decoding performance was related to the number of regions included in the model, we used whole-spectrum power, coherence, and directionality within all possible combinations of specified numbers of regions (from 1 to 7).

#### Decoding by frequency

To evaluate the decoding performance of specific frequencies, instead of fitting the LM against the whole spectrum (50 bins), we fitted it against each frequency bin (1 bin). Because the number of variables for each bin (1 to 7 for power, 1 to 21 for coherence and 1 to 21 for directionality) was, sometimes close to the number of PCs used for other analyses (20), we calculated a curve for every cumulative number of PCs from 1 to 20 (PC1, PC1 to PC2, PC1 to Nth PC, until PC1 to PC20). For region × frequency decoding, we fitted the GLM on UV data because it had only 13 attributes (1 power, 6 coherence and 6 directionality). We compared the rats’ average AUC across cumulative PCs between frequency ranges (theta: 4-10 Hz; no-theta: 1-4 plus 10-50 Hz) and the proportions of maximum AUC within frequency ranges across all PCs of all rats (N = 120).

#### Decoding and CS association

We examined the dependence of decoding performance on the association between CS to either fear or safety by training the models to discriminate CSs between two sessions for every pair of days. Then, we compared the average performances between CS+ vs. CS-, CS+ vs. CS+ and CS-vs. CS-.

#### Generalization

To estimate within-subject generalization (test performances) of decoder models, we trained a model on data from the Nth day (1st, 2nd, 3rd, and on) of safety versus the Nth day of the fear protocol and performed out-of-sample testing on every remaining Nth pair of days. Then, we obtained the average of test performances for all Nth days trained and tested. For AA, we trained the model on each session and tested it on all others, irrespective of the protocol. We also compared the performances with the GLM fitted on UV data, which we expected to exhibit overfitting. To estimate between-subject generalization, we trained the model on all epochs of a single subject and averaged the test performances on all other subjects.

#### Artificial neural network

We also used a shallow artificial neural network (ANN) consisting of a single hidden layer with 20 neurons with sigmoid transfer function and 1 output neuron with softmax transfer function. Each ANN was trained using symmetric random weight/bias initialization and updates according to the scaled gradient conjugate method, 70/30% train/test data divided randomly, and cross-entropy performance error estimate. Training would stop after 6 subsequent iterations with increases of validation error. We trained the ANN 10 times and selected the one with the highest performance. The ANN were trained only on UV data.

#### Individual variability

To visualize the dispersion of spectral data across individuals and stress-related states, we used two-dimensional t-distribution stochastic neighbor embedding (t-SNE) of the first 20 PCs of Z-scored data (across subjects) with the following parameters: barneshut algorithm with 0.5 tradeoff parameter, Euclidean distance, exaggeration size of 4, perplexity of 30 local neighbors, and normally distributed random (×10^-4^) initialization. We clustered t-SNE data with k-means using squared Euclidean distances, 1000 replicates, and 1000 maximum number of iterations. To estimate within- and between-subject variance, we computed the mean squared Euclidean distance of the first 20 PCs of Z-scored data across all epochs of interest of each subject and the average between each rat to all others, respectively.

#### Electome analysis

We Z-scored each subject prior to model training on all rats for decoding states from all individuals simultaneously (electome analysis). The dot product between GLM coefficients (for each PC) and PC coefficients (for each variable) returned the LM coefficients for each variable (region/pair × frequency bin × estimate), reflecting its weight and contribution to the model.

### Statistical analyses

Normality was assessed using the Lilliefors test. We used paired Student t-test for within-subject comparisons and multiple t-tests for frequency-wise comparisons. We used repeated measures (RM) one-way analysis of variance (ANOVA) for comparisons between features or models and RM two-way ANOVA for two features or across time (days or seconds), and Fisher’s least significant difference (LSD) test as *post hoc* analysis. We performed mixed-effects models using the restricted maximum likelihood method for RM ANOVA with missing values. We examined the trend relationship of variables with time or number of regions using linear or nonlinear (x as log and y as linear) regression and F-test for nonzero slope. The chi-squared test was used to compare the proportions of maximum AUC within and out of theta range. We calculated Pearson’s correlation between normal distributions and Spearman’s rank correlation as a non-parametric equivalent. Data are expressed as the median, second-third quartiles range, and multiple significances are expressed as the minimum-maximum range. The significance level was set to 0.05 unless stated otherwise.

## Results

### Differential conditioning to safety or fear promotes approach or avoidance in a within-subject design

To investigate how multiple brain structures encode fear or safety and approach (AP) or avoidance (AV) we recorded LFP from seven regions of the mesolimbic-hippocampal-frontocortical network (MLHFC; Fig. 2*B-C*) in a newly developed protocol that delineates such behavioral states. Here, a conditioned stimulus (CS) was associated with safety (CS-; absence of shocks) in the initial days and then to fear (CS+; anticipation of shocks), and these associations were tested in an approach-avoidance (AA) test daily (Fig. 1*A-D*).

Initially, we confirmed that all rats performed the escape response to the US throughout sessions irrespective of CS association (mixed-effects RM one-way ANOVA: session F(9,37) = 1.48, p = 0.18; Fig. 1*E*). In turn, we observed greater probabilities of avoidance attempts out of CS- and during CS+ throughout sessions (Fig. 1*F*) and on average (t(5) = 18.67, p < 0.0001), indicating the differential association with safety and fear, respectively. In the AA test, we expected that CS-conditioning would increase the probabilities of approach and time spent on reward during CS, while the CS+ would promote the opposite effect and increase the probabilities of avoidance and time on platform (Fig. 1*D*). Indeed, we observed a learning curve of the probabilities of approach *versus* avoidance from the start of safety conditioning and after reversal to fear (mixed-effects RM two-way ANOVA: session × behavior interaction F(9,32) = 8.78, p < 0.0001; Fig. 1*G*). In accordance, we only found significant differences between probabilities of AP vs. AV after the second day, which remained until the last safety session (t(36) = 2.65-4.59, p = 5.1×10^-5^-0.01; Fig. 1*G*). In turn, there was no difference on the following days after reversal except for the third fear session onwards (t(df) = 2.29-3.83, p = 0.0004-0.02; Fig. 1*G*). A similar pattern was found for the probabilities of time spent in reward versus on platform (RM mixed-effects model: session × behavior interaction F(9,32) = 5.50, p = 0.0001), as the probability of time on reward was significantly greater after the second day until the last safety session (t(36) = 2.77-4.49, p = 7.0×10^-5^-0.008; Fig. 1*H*). Taking the average of all sessions, there were significant increases in the probabilities of avoidance (t(4) = 3.57, p = 0.02) and time on platform (t(4) = 5.80, p = 0.004) during CS+ than CS-. Although there were no significant differences on average, the probabilities of approach (t(4) = 4.69, p = 0.009) and time on reward (t(4) = 2.35, p = 0.07) during CS were greater on the last session of safety than fear (Fig. 1*G-H*).

In conclusion, this new experimental design was able to promote a differential conditioning to safety and fear, as expressed by the probabilities of approach and avoidance during CS on test sessions. Thus, this proves to be an appropriate behavioral protocol to investigate neural activities and within-subject decoding of fear, safety, approach and avoidance states.

### Increased mesolimbic-hippocampal-prefrontal cortical theta activity in fear and approach

Previous studies indicated the involvement of theta oscillations in processing fear and anxiety. Before investigating the multivariate decoder models, we examined the participation of theta activity during fear versus safety and in the prediction of approach or avoidance through common univariate analysis.

Notably, we observed a clear differentiation between stress-related states within the theta range (Fig. 3). For CS+/CS-, it preferentially occurred within 5-9 Hz (Fig. 3*A*) and, for AA, within 8-11 Hz (Fig. 3*B*). By frequency-wise comparisons, we found that CS+ was associated with increased theta coherence (t(4) = 2.24-5.94, p = 5.10×10^-6^-0.049) in a tegmental-hippocampal-frontocortical subnetwork (PL-[DH, VH, VTA]; IL-[DH, VH, VTA]) with lower values between VTA-NAc and DH-AMG (Fig. 3*C*). In addition, we found changes in directionality (t(4) = 2.34-11.95, p = 3.01×10^-7^-0.049) in a limbic-frontocortical subnetwork (PL-[NAc, AMG], IL-[NAc, AMG]) rather than coherence (Fig. 3*C*).

**Figure 3.**
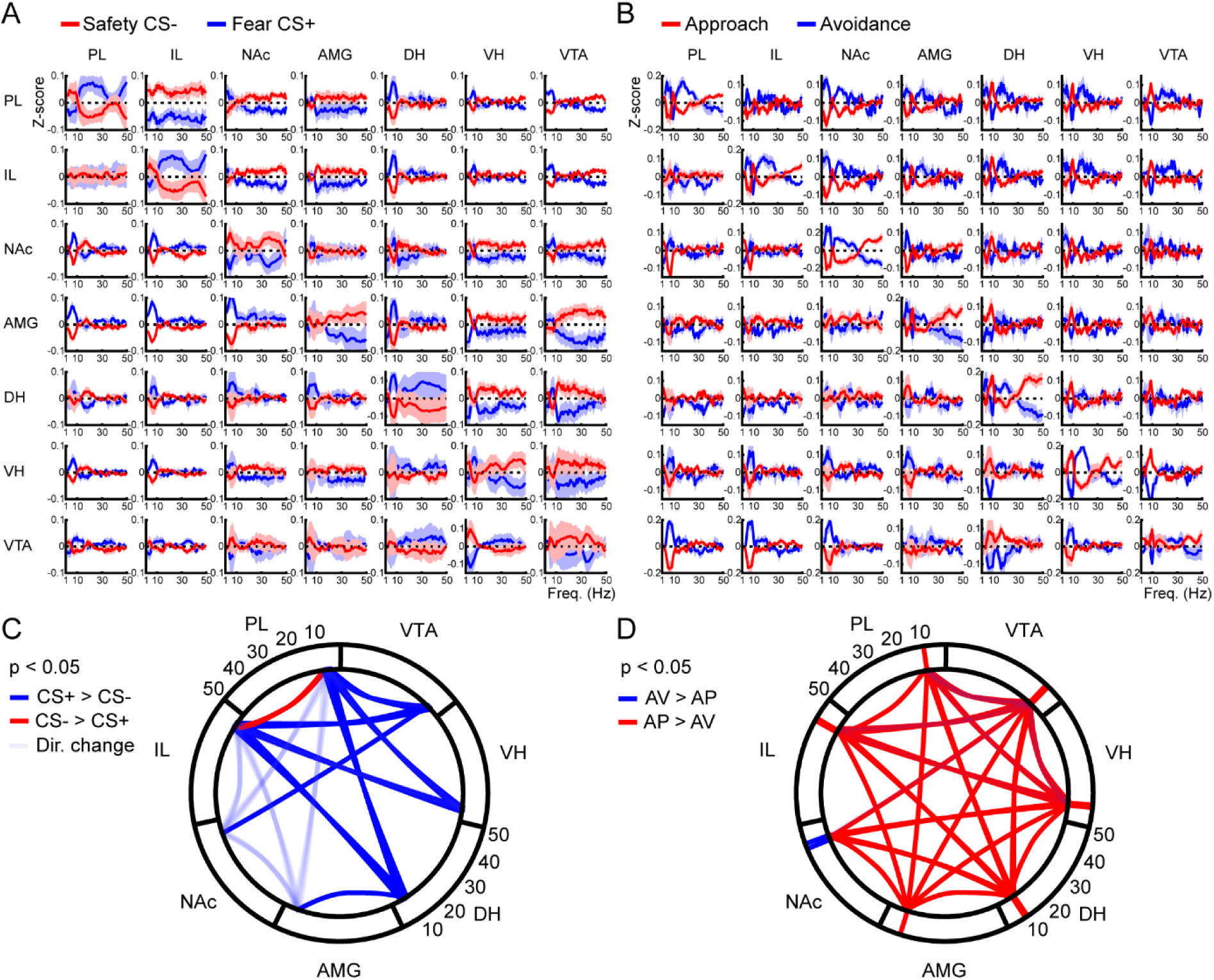
Increased mesolimbic-hippocampal-prefrontal cortical theta activity in fear and approach. ***A-B,*** Average comparisons of Z-score transformed spectral estimates. Diagonal panels show power, upper right panels show coherence, and lower left panels show directionality (phase difference between y-axis region minus x-axis region) comparisons. ***C-D*,** Significant differences (t-tests p < 0.05) within theta ranges for **(*C*)** CS+/CS-(4-9 Hz) and **(*D*)** approach (AP) and avoidance (AV) (8-11 Hz).

In the AA sessions, we found a remarkable increase in theta power and coherence on the periods preceding AP, compared to AV (Fig. 3*B*). These increases were significant (power: t(4) = 2.40-5.49, p = 2.70 × 10^-5^-0.049; coherence: t(4) = 2.35-9.80, p = 6.11×10^-7^-0.05) for almost all regions and pairs of regions (except PL- and IL-NAc; Fig. 3*D*). In addition, we found changes in directionality in the tegmental-hippocampal-frontocortical subnetwork that differentiated AP and AV (t(4) = 2.34-5.22, p = 4.12×10^-5^-0.049; Fig. 3*D*). Surprisingly, only the NAc showed greater power in the periods preceding AV, but looking at the spectrum (Fig. 3*B*, *D*), this difference was due to overall effects on the whole spectrum rather than specifically in theta. In fact, we also noticed some differences beyond theta in some pairs of regions (Fig. 3*B*), suggesting activities in other frequencies may also be involved.

Our findings corroborate previous evidence for the association between increased limbic-cortical theta power and coherence in fear and directionality changes associated with safety. However, we found stronger theta power and synchrony preceding approach rather than avoidance. In addition, we showed that the relationship between theta activity and stress coping happens at multiple regions comprising a brain-wide network.

### Brain-wide power, synchrony and directionality decode fear or safety better than each region separately

To explore how different sets of features of oscillatory activity can decode fear or safety, we developed a linear decoding pipeline. Importantly, our behavioral protocol and signal quality criteria were able to provide a proper (CS- = 2066, 1877-2979; CS+ = 1994, 1247-2596; AP = 457, 425-1341; AV = 457, 350-1076) and balanced (CS+/CS- = 48.99%, 43.36-51.35%; AV/AP = 45.25%, 42.10-47.89%) number of epochs per rat for classification. First, we addressed the issue that by choosing different features, the number of attributes/variables would change, which directly affects the performance of classification models. For instance, taking the whole spectrum (50 frequency bins) of all regions (7 regions, 21 pairs) and estimates (power, coherence, direction) would comprise 2450 attributes (Fig. 2*E*), while taking the power of a single region would contain only 50 attributes. Therefore, we performed dimensionality reduction to a fixed number of multivariate scores and fitted a generalized linear model.

We investigated several dimensionality reduction methods (e.g., factor analysis, hierarchical clustering, non-negative matrix factorization) and chose PCA because it is one of the most widely studied, faster and interpretable methods. PCs are linear combinations of correlated data. Therefore, the PCs from our data reveal collective variations of spectral estimates across regions and frequencies, as illustrated by representative PCs’ coefficients of power and coherence (Fig. 2*F*), indicating that these scores represent network activities. In contrast, PCs are uncorrelated to each other, avoiding issues with multicollinearity. Moreover, PCs are ordered by the amount of variance they explain, and so the number of PCs for proper decoding can also reveal biological information.

We found that, on average, the best PC (from all estimates combined) had a greater AUC than the best UV variable (across all frequencies, regions and pairs; RM one-way ANOVA F(3,15) = 4.70, p = 0.01; Fig. 2*G*) for power (t(15) = 2.04, p = 0.058), coherence (t(15) = 3.71, p = 0.002), and directionality (t(15) = 2.34, p = 0.03), confirming that these multivariate scores contain relevant information for stress-related decoding. Then, we observed that most of the relevant (AUC > 0.60) PCs (for power, coherence, directionality, and all estimates combined) for discriminating fear vs. safety occurred until the 20th PC (Fig. 2*H*), so we heuristically determined this number to evaluate decoding performances throughout this study. Importantly, the number of epochs for each rat was several times greater than this number of PC scores used for GLM regression, which also aided in avoiding overfitting. In fact, we observed that GLM coefficients closely corresponded to each PC AUC (see example in Fig. 2*I*), also suggesting a lack of overfitting.

Having established a decoding pipeline that avoids the bias of the number of attributes, we set out to investigate how the activity of different sets of multiple brain structures and spectral estimates can decode fear or safety. First, we investigated how each region and estimate (local power, coherence, and directionality to all other regions) would classify fear or safety. We found that every region and estimate showed a greater AUC than the models trained on shuffled labels (t(5) = 2.97-12.26, p = 6.37×10^-5^-0.03; not shown). Then, we also computed the model considering all regions simultaneously, and found that the decoding performance, considering each estimate and all estimates combined, was greater than when each region was considered separately (RM two-way ANOVA region F(7,35) = 3.84, p = 0.003; *post hoc*: t(35) = 2-19-4.14, p = 0.0002-0.003; Fig. 4*A*, *C*). We also found a significant interaction between region and estimate (RM two-way ANOVA region × estimate interaction F(21,105) = 2.00, p = 0.01; Fig. 4*C*). In particular, we noticed that the DH showed greater performance than other regions (Fig. 4*A*, *C*). Indeed, all regions-all estimates models showed significant difference to every separate region (t(105) = 4.05-5.31, p < 0.0001), but only a tendency against the DH (t(105) = 1.95, p = 0.053).

**Figure 4.**
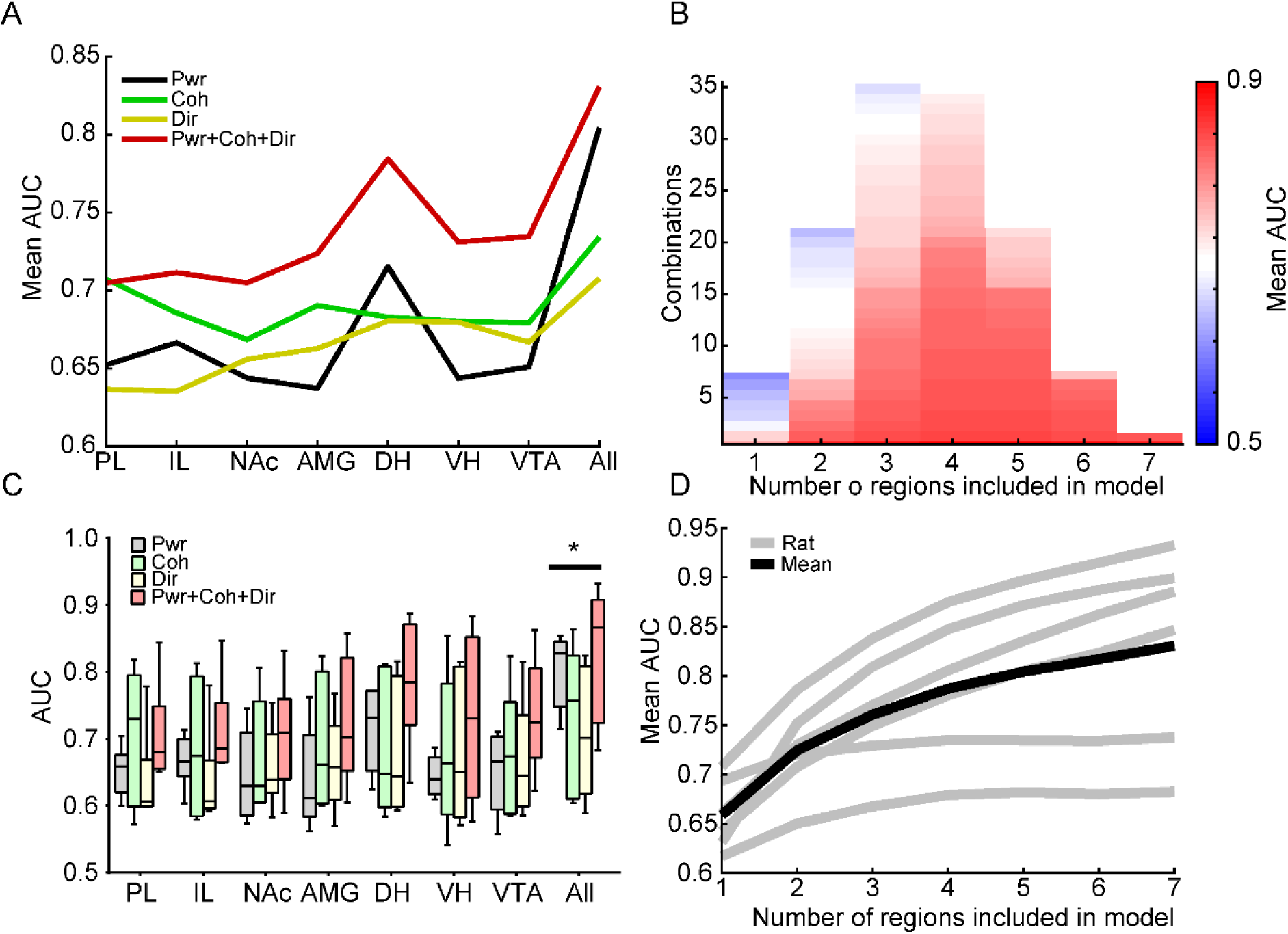
Brain-wide power, synchrony, and directionality decode fear or safety better than each region separately. ***A, C*,** AUC of each region and respective estimates show better performance if all regions and estimates are considered together. *p < 0.05 (Fisher’s LSD test for comparisons between regions; all vs. every region). Note the particularly good decoding of the DH. ***B,*** Ranked AUC from all possible combinations of regions in ascending order. ***D*,** Average decoding performance increases as a function of the number of regions included. This pattern is similar for all rats. Here and on: box plots represent the median and interquartile range, and whiskers represent the minimum and maximum.

Next, we investigated the relationship between decoding performance and the number of regions included in the model. We calculated the AUC of models from all possible combinations of regions and their power-coherence-directionality values (Fig. 4*B*). We found that the average decoding performance increased as a function of the number of regions included in the model (number of regions × mean AUC linear regression: r(5)^2^ = 0.90, F(1,5) = 47.31, p = 9.93×10^-4^; nonlinear regression: r(5)^2^ = 0.99, F(1,5) = 82207, p < 1×10^-10^), reaching the highest performance if all regions were included (Fig. 4*D*). Remarkably, all rats showed this same pattern (linear regression: r(5)^2^ = 0.67-0.96, F(1,5) = 10.21-138.73, p = 7.76×10^-5^-0.01-0.02; nonlinear regression: r(5)^2^ = 0.88-0.99, F(1,5) = 3630-75252, p = 1×10^-10^-2.38×10^-8^; Fig. 4*D*).

We conclude that the best classification performances are achieved if all regions and estimates are included for the model generation, although some regions and estimates may show greater relevance for decoding fear or safety, such as the DH. Our results evidence that fear and safety are encoded at the brain-wide level, as represented by patterns of collective variations across multiple regions simultaneously.

### Patterns of brain-wide theta activity decode fear or safety better than other frequencies

After showing that whole-spectrum data can decode fear or safety, we sought to investigate what frequencies were the most relevant. We estimated the decoding performance for every frequency across cumulative numbers of PCs. Because PCs are ranked by the proportion of data variance they explain, we hypothesized that the frequencies where variation was more related to fear vs. safety would exhibit good performance already on more initial PCs. From the average PC × frequency classification, we noticed markedly greater performances within 4-10 Hz, which started already at the first 5 PCs (Fig. 5*A*). Indeed, the averaged performances within theta were significantly greater than non-theta frequencies (t(5) = 4.32, p = 0.007; Fig. 5*B-C*) and the proportion of maximum AUCs within theta (N = 94/120, 78.33%) was significantly greater than out of theta (X^2^(1, N = 120) = 77.07, p < 0.0001; Fig. 5*D*). Moreover, we observed that theta frequencies showed average greater performances for all estimates (Fig. 5*A*), power, coherence, directionality, and every region (Fig. 5*E*).

**Figure 5.**
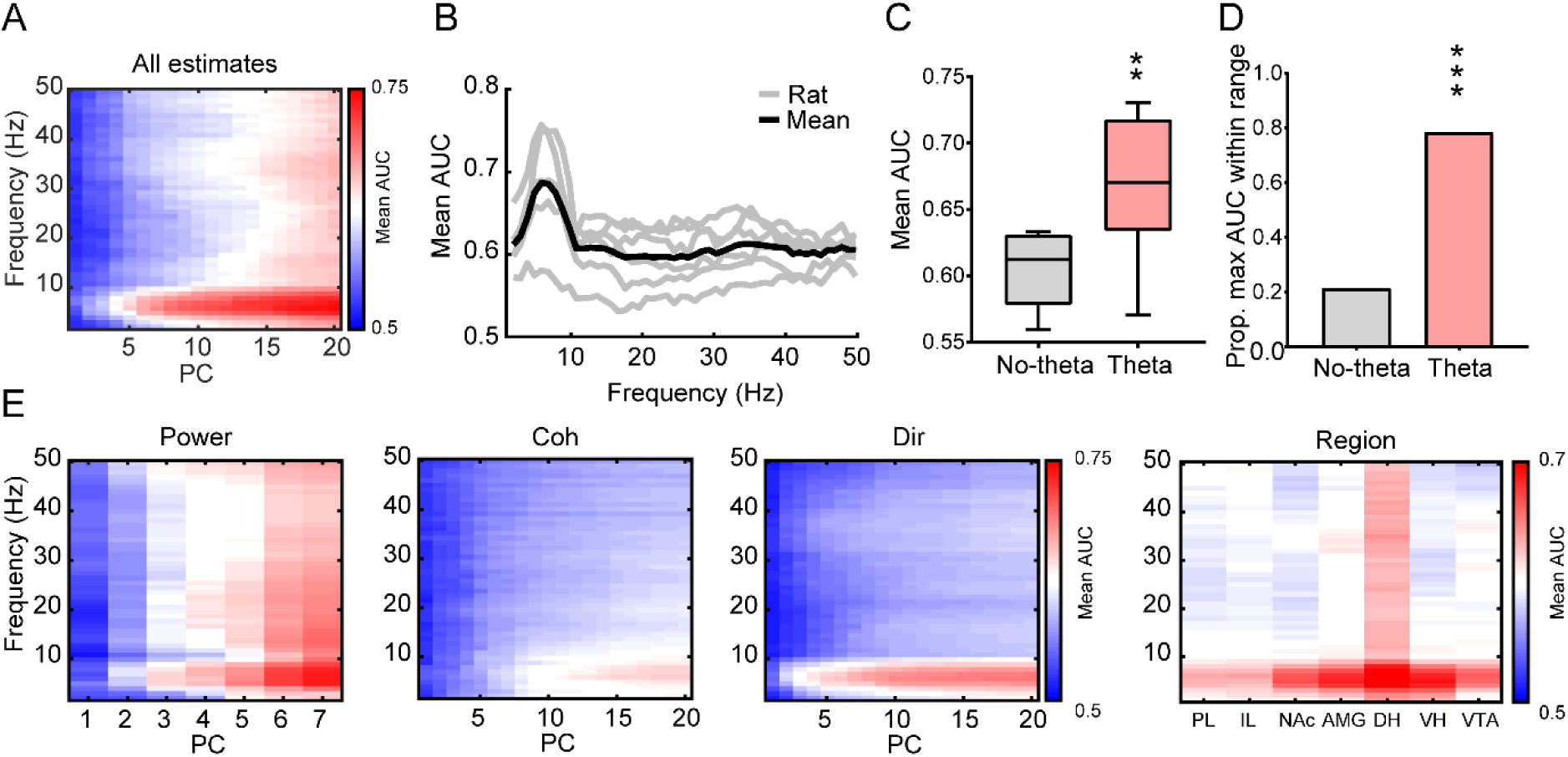
Patterns of brain-wide theta activity decode fear or safety better than other frequencies. ***A-D*,** Brain-wide power, coherence, and directionality collectively shows greater decoding performances within the theta frequencies. Note good decoding in the theta range with fewer PCs. Theta frequencies show significantly greater **(*C*)** AUC (paired t-test **p < 0.01) and **(*D*)** proportion of maximum AUC across all rats and PCs (chi-squared test ***p < 0.0001) than other frequencies. ***E*,** The better decoding of theta frequencies happens separately for power, coherence, directionality, and every region.

In sum, we show that theta network activity is the most relevant for decoding fear or safety. These findings corroborate previous reports, but demonstrate that all regions and estimates are relevant for this performance, indicating a complex network activity involving amplitude, synchrony, directionality and multiple regions simultaneously. Furthermore, such a preference of the decoder model to these specific frequencies validates the physiological relevance of the algorithm used.

### Decoding CS association to safety or fear predicts approach or avoidance

To investigate if decoder performances were associated with CS association to fear or safety, first, we examined the model performances trained between CSs from all sessions. We observed that the performances were better when decoding CS+ vs. CS-(RM one-way ANOVA F(2,10) = 9.76, p = 0.004) than CS-vs. CS-(t(10) = 3.12, p = 0.01) or CS+ vs. CS+ (t(10) = 4.26, p = 0.001; Fig. 6*A*). This finding demonstrates a greater dependence of decoding performance to CS association than to inherent variability across days. Then, we noticed that decoder scores, trained on all data of each individual, did not return a homogeneous distribution across sessions, but rather, it exhibited a temporal effect (mixed-effects RM one-way ANOVA F(9,37) = 10.27, p < 0.0001; Fig. 6*B*). We observed from the averaged scores a poor decoding performance in the first sessions of safety and fear, which was similar to the learning curve for behavioral measures of fear and safety described earlier (Fig. 6*B*; see also Fig. 1*G-H*). Therefore, we hypothesized that the decoder scores could be correlated to the behavioral measures of CS association, evaluated later on test sessions. Indeed, decoder scores (positive values for CS+) during conditioning showed positive correlations to the probabilities of avoidance (r(45) = 0.56, p < 0.0001) and time on platform (r(45) = 0.44, p = 0.001), while showing negative correlations with the probabilities of approach (r(45) = −0.33, p = 0.01) and time on reward (r(45) = −0.35, p = 0.01) during test sessions (Fig. 6*C*). Thus, we conclude that CS+/CS-brain-wide decoding is dependent on the association between the CS to either fear or safety, as the decoder scores predicted behavioral manifestations of these associations.

**Figure 6.**
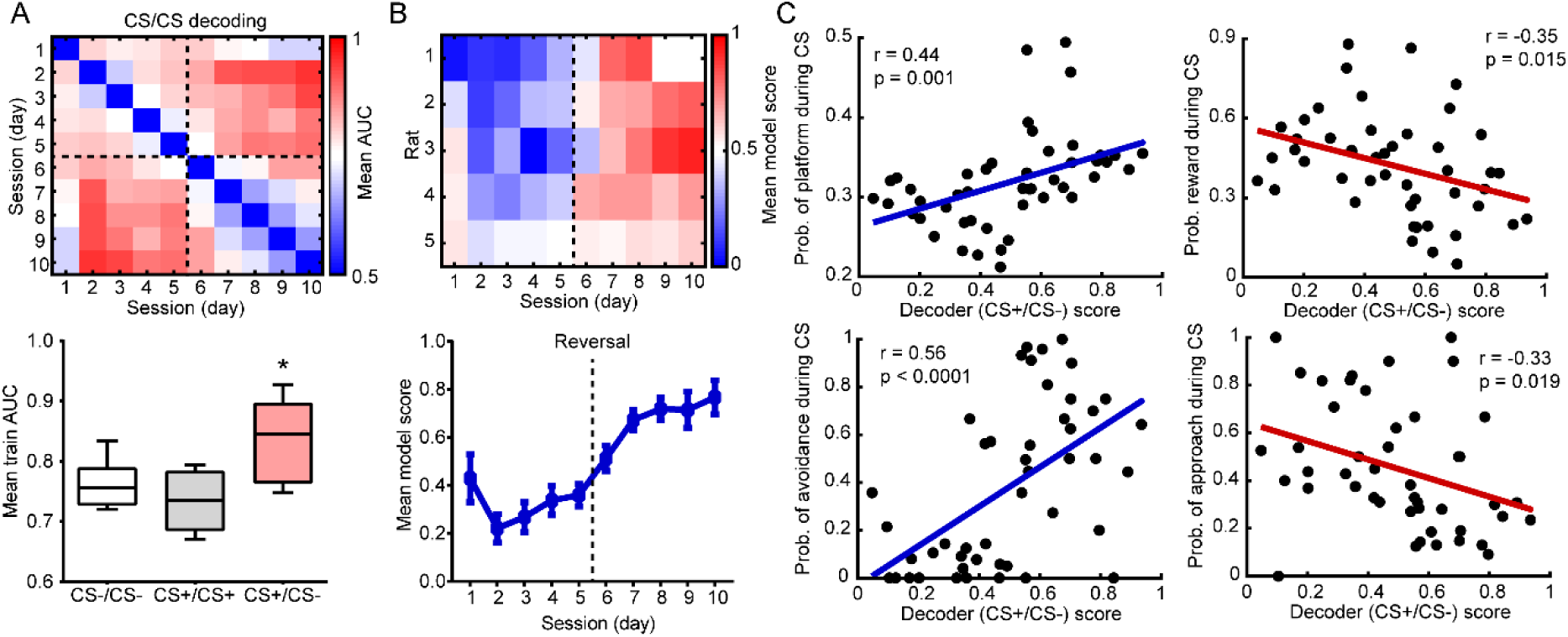
Decoding CS association to safety or fear predicts approach or avoidance. ***A*,** Decoding performances across days depended on CS association. The average AUC trained between days was greater for CS-vs. CS+ than for CS+ vs. CS+ or CS-vs. CS-. *p < 0.05 (Fisher’s LSD test). ***B*,** Heterogeneous distribution of decoder scores throughout sessions. Average decoder scores resemble the learning curves of behavioral measures of fear and safety (see Figure 1). ***C*,** Average decoder scores of CS (CS+ = 1, CS-= 0) during conditioning positively correlated with the probabilities of avoidance and time on platform and negatively correlated with the probabilities of approach and time on reward during CS on test sessions.

### Within-subject generalization of decoder models for fear or safety

We investigated if decoder models exhibit generalization, that is, whether they perform well when tested on new unseen data. Remarkably, we observed that decoder models showed good average performances when tested on sessions they were not trained on (AUC = 0.76, 0.66-0.90; Fig. 7*A*). In turn, the LM trained on UV (LMUV) exhibited overfitting, as they showed excellent train (AUC = 1) but bad test (AUC = 0.54, 0.52-0.59) performances (Fig. 7*C*), likely because the numbers of variables and observations were close in this case. In fact, the LM trained on PCs (LMPC) showed greater test performances than the LMUV (t(4) = 4.58, p = 0.01; Fig. 7*C*). We also trained an ANN on UV, which is usually considered a gold standard classification model, and we found an equivalent generalization to the LMPC (AUC = 0.76, 0.68-0.89; t(4) = 0.30, p = 0.77; Fig. 7*C*). Next, we investigated between-subject generalization by training the model on one rat and testing on all others (Fig. 7*B*). Interestingly, we found a poor between-subject generalization for both LMPC (AUC = 0.48, 0.47-0.53) and ANN (AUC = 0.50, 0.49-0.53; Fig. 7*B, C*). In addition, the test performances were significantly lower than within-subject performances (LMPC: t(9) = 5.11, p = 0.0006; ANN: t(9) = 5.27, p = 0.0005 Fig. 7*C*). The better performance of within-subject compared to between-subject generalization reveals a relevant individual variability in stress processing. Also, the generalization of the LM models and the equivalent performance to ANN substantiate the validation of the decoding method.

**Figure 7.**
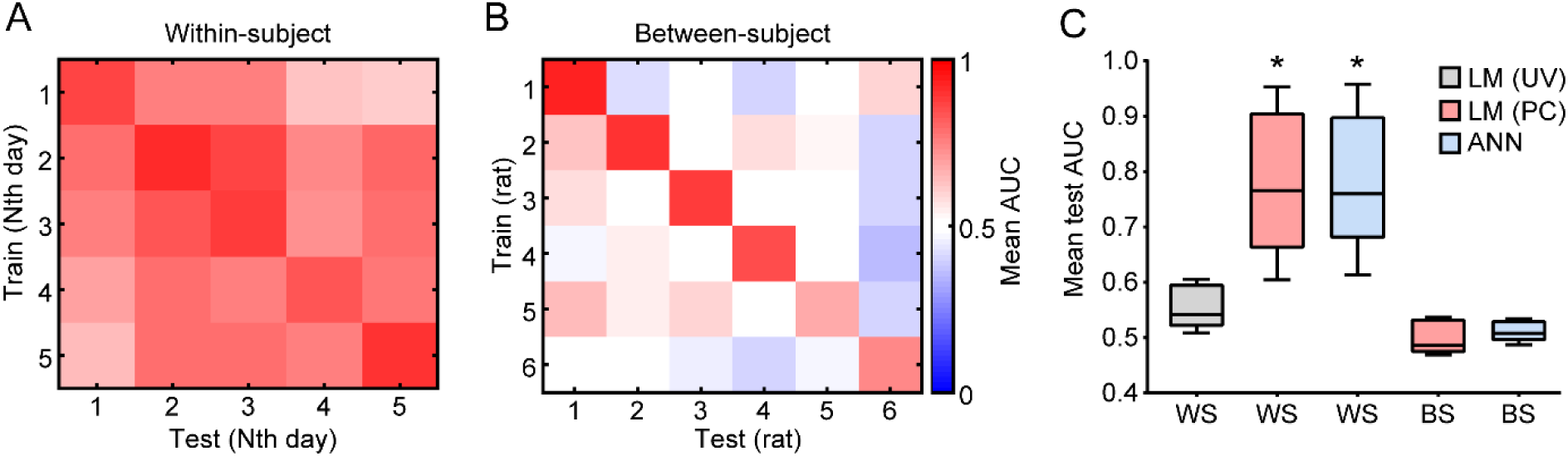
Within-subject generalization of decoder models for fear or safety. ***A*, *B*,** Decoder models exhibited **(*A*)** robust within-subject generalization across days but **(*B*)** poor generalization between rats. ***C*,** The linear model (LM) trained on univariate (UV) data exhibited overfitting, while trained on PC presented a good generalization equivalent to an artificial neural network (ANN). Both LM and ANN showed poor between-subject validation. WS = within-subject; BS = between-subject. *p < 0.05 (paired t-test against LMUV).

### Individual variability of brain synchrony patterns unrelated to stress

After observing such a diversity of individual variability in stress processing, we investigated how the patterns of brain-wide activity differ between individuals. We observed that there is an interindividual variability of brain power and synchrony, irrespective of stress-related states (Fig. 8*A*). t-SNE transform of brain-wide activity returned six well-defined clusters (Fig. 8*B*), which precisely corresponded to each individual rat (k-means clusters × rats: N = 25368/25399 epochs, 99.88%). We also noticed from t-SNE inspection that interindividual variability is much more relevant than stress-related information, as CS+/CS-epochs showed significant overlap within each rat cluster (Fig. 8*B*). We assessed further the within-and between-subject variability, measured by the mean squared distances and we confirmed a significantly greater between-subject than within-subject variance for power (t(5) = 3.88, p = 0.01), coherence (t(5) = 9.32, p = 0.0002), directionality (t(5) = 3.74, p = 0.01) and all estimates combined (t(5) = 6.67, p = 0.001; Fig. 8*C*). In addition, we found that coherence is the measure with the greatest interindividual variability. Therefore, our findings indicate a substantial individual variability in both stress coping-related processing and invariant brain activity.

**Figure 8.**
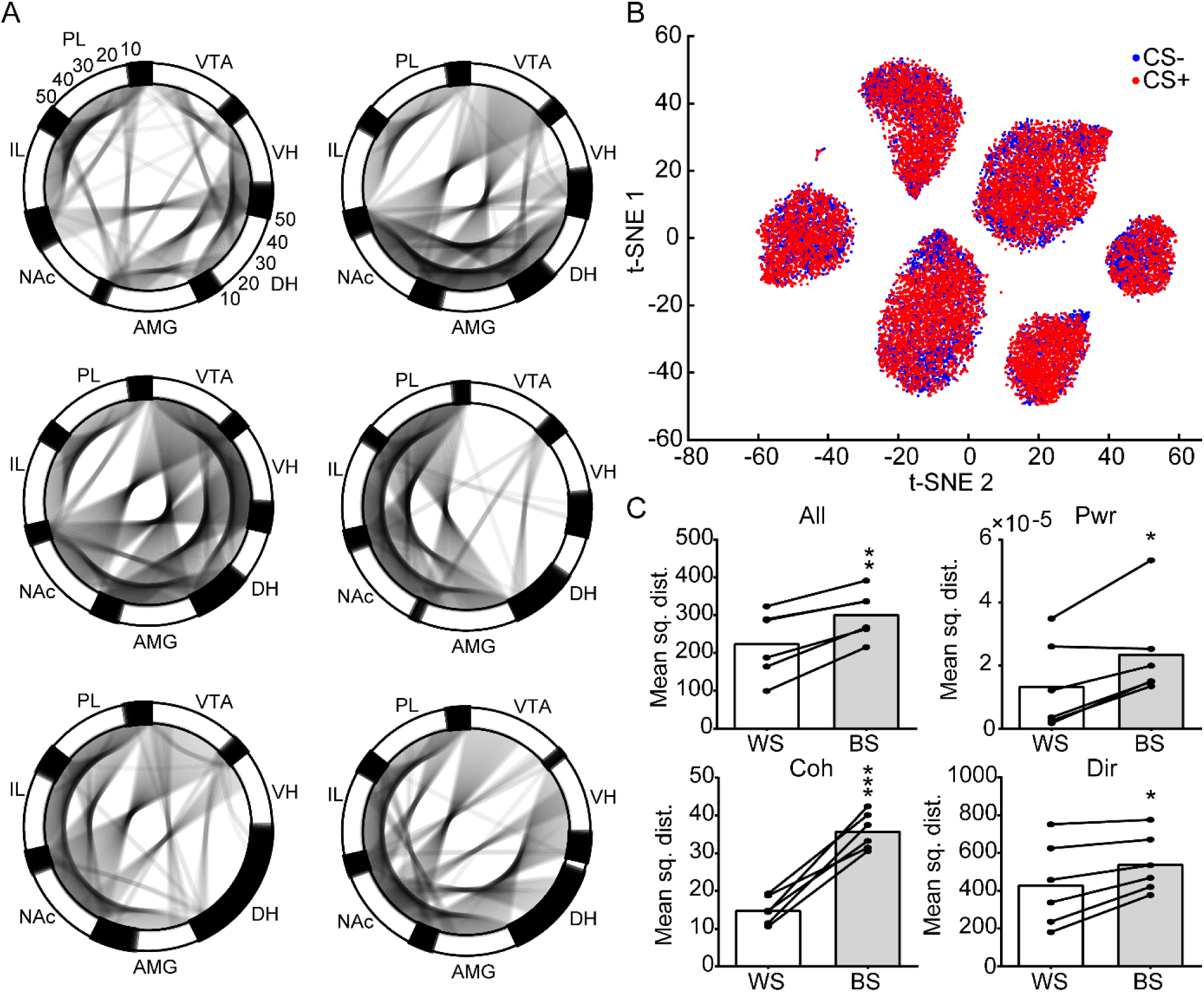
Individual variability of brain synchrony patterns unrelated to stress. ***A*,** Network chord diagrams representing the average power and coherence estimates for each rat. Note the variability of whole-spectrum coherence strengths between pairs of regions across rats. Note that brain-wide synchrony resembles a small-world network configuration. ***B,*** Two-dimensional t-distributed stochastic neighbor embedding (t-SNE) of spectral data showing six well-defined clusters that correspond to the six rats, while CS+ and CS-clearly overlap. ***C,*** All estimates show greater average mean squared distances between subjects than within each. Note the more significant variances between subjects for coherence estimates. WS = within-subject; BS = between-subject. *p < 0.05, **p < 0.01, ***p < 0.001 (paired t-test).

### Decoding approach versus avoidance reveals similar preferences for brain-wide and theta activities, and within-subject generalization

After validating the decoding pipeline, we sought to investigate how the same method could be applied to decode AP and AV motivational states. Strikingly, AA decoding revealed similar characteristics to CS+/CS-decoding (Fig. 9). We observed greater performance from all estimates if all regions are considered rather than each one separately (RM two-way ANOVA region F(7,28) = 5.11, p = 0.0007; t(28) = 2.19-5.02, p = 2.6×10^-5^-0.03; Fig. 9*A*, *D*). We also observed a growing curve of the average performance as function of the number of regions included in the model (linear regression: r^2^(5) = 0.90, F(1,5) = 49.23, p = 9.06×10^-4^; nonlinear regression: r^2^(5) = 0.98, F(1,5) = 46203, p = 4.13×10^-11^; Fig. 9*B*, *E*). However, differently from CS+/CS-, the NAc showed the greatest performance, particularly for power data, which is also consistent with the univariate results (Fig. 9*A*, *D*; see also Fig. 3*B*). Also in agreement with the univariate analysis, the theta frequencies showed the greatest performances across the spectrum (t(4) = 4.40, p = 0.01; Fig. 9*C*, *F*). Interestingly, we observed a better performance for within-subject generalization (AUC = 0.66, 0.59-0.76; Fig. 9*G*), which was significantly greater than the LMUV overfitted model (t(4) = 4.20, p = 0.01; Fig. 9*I*), but was not significantly different from between-subject classification (t(4) = 1.74, p = 0.15; AUC = 0.60, 0.53-0.65; Fig. 9*I*). In fact, AA decoding showed a slightly greater between-subject validation than CS+/CS-(Fig. 9*H-I*; see also Fig. 7*B*, *D*). In conclusion, our decoding pipeline exhibits generalized applicability, as it can decode distinct dimensions of stress coping. Also, both AA and CS+/CS-decoding showed similar characteristics, supporting the preference for brain-wide, theta frequencies, and individual variability in the encoding of stress coping-related information.

**Figure 9.**
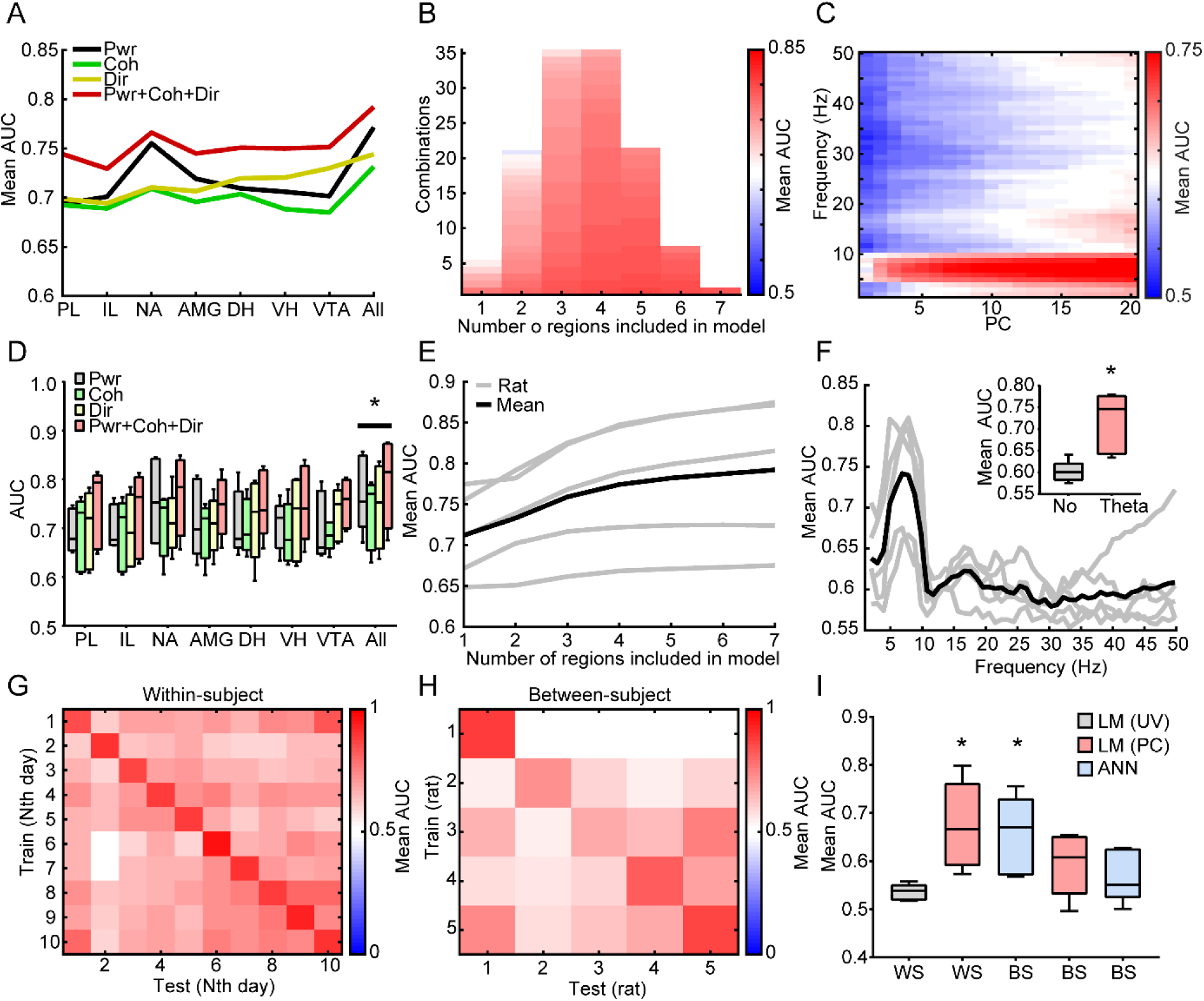
Decoding approach versus avoidance reveals similar properties to decoding fear versus safety. ***A*, *C*,** All regions and estimates combined showed greater decoding than each separately. *p < 0.05 (Fisher’s LSD test for comparisons between regions; all vs. every region). Note particular good decoding of the NAc. ***B*,** Ranked AUC from all possible combinations of regions in ascending order. ***E*,** Average decoding performance increases as a function of the number of regions included in all rats. ***C*, *F*,** Specifically great decoding within the theta frequencies occurred with **(*C*)** fewer PCs and **(*F*)** in all rats. *p < 0.05 (paired t-test). ***G-H*,** Decoder models exhibited greater within-subject generalization across days than between rats. However, note a moderate performance of between-subject validation. ***I*,** The linear model (LM) trained on univariate (UV) data overfitted while trained on PC showed a good generalization equivalent to an artificial neural network (ANN). Both LM and ANN showed moderate between-subject generalizations. *p < 0.05 (paired t-test against LMUV).

### Patterns of brain-wide activity decode fear or safety and approach or avoidance across individuals

Although we found a poor between-subject generalization of decoder models if they were trained in specific individuals, we sought to investigate if our decoding pipeline could also identify patterns that decode stress coping-related states across subjects. Despite using a different algorithm, our ultimate results are equivalent in terms of spectral estimates and interpretation to that of Hultman et al. (2018), so we will use the same nomenclature to address these patterns of electrical functional connectomes as electomes.

Remarkably, the most prominent characteristics of both the fear electome and the approach electome were brain-wide power and synchrony in the theta range (Fig. 10*A-B*). Particularly in the fear electome, it was observable a slight prominence within the beta (12-30 Hz) band (Fig. 10*A*). Taking the average of electome scores for all CS+ and CS-periods and pre-AP and pre-AV, the electomes identified correctly all stress-related states from all subjects (CS+ vs. CS-t(4) = 2.86, p = 0.04; AUC = 100%; Fig. 10*C*; AP vs. AV t(4) = 4.33, p = 0.01; AUC = 100%; Fig. 10*E*). Similar to within-subject decoding, CS+/CS-decoder scores significantly correlated with later probabilities of avoidance on the test sessions (r(45) = 0.48, p = 0.0005; Fig. 10*D*), showing that the decoding dynamics were related to the learning dynamics. Hence, we hypothesized that AA decoder scores throughout the epochs could also reveal biologically relevant information. For that, we trained the model considering 10 s before until 1 s after each crossing. We observed that the approach electome score gradually increased prior to the crossing, so the strength of this activity predicted the behavior (time × average score linear regression: r^2^(9) = 0.81, F(1,9) = 39.15, p = 0.0001; Fig. 10*F*). However, by looking at the pre-AV periods we saw a deeply lower electome score that gradually increased before AV crossings too (linear regression: r^2^(9) = 0.90, F(1,9) = 82.86, p = 7.77×10^-6^; Fig. 10*F*). Interestingly, the electome scores converged prior to both AP and AV crossings (RM two-way ANOVA time × valence interaction F(10,40) = 2.93, p = 0.007), suggesting that this activity may underlie both motivational valences.

**Figure 10.**
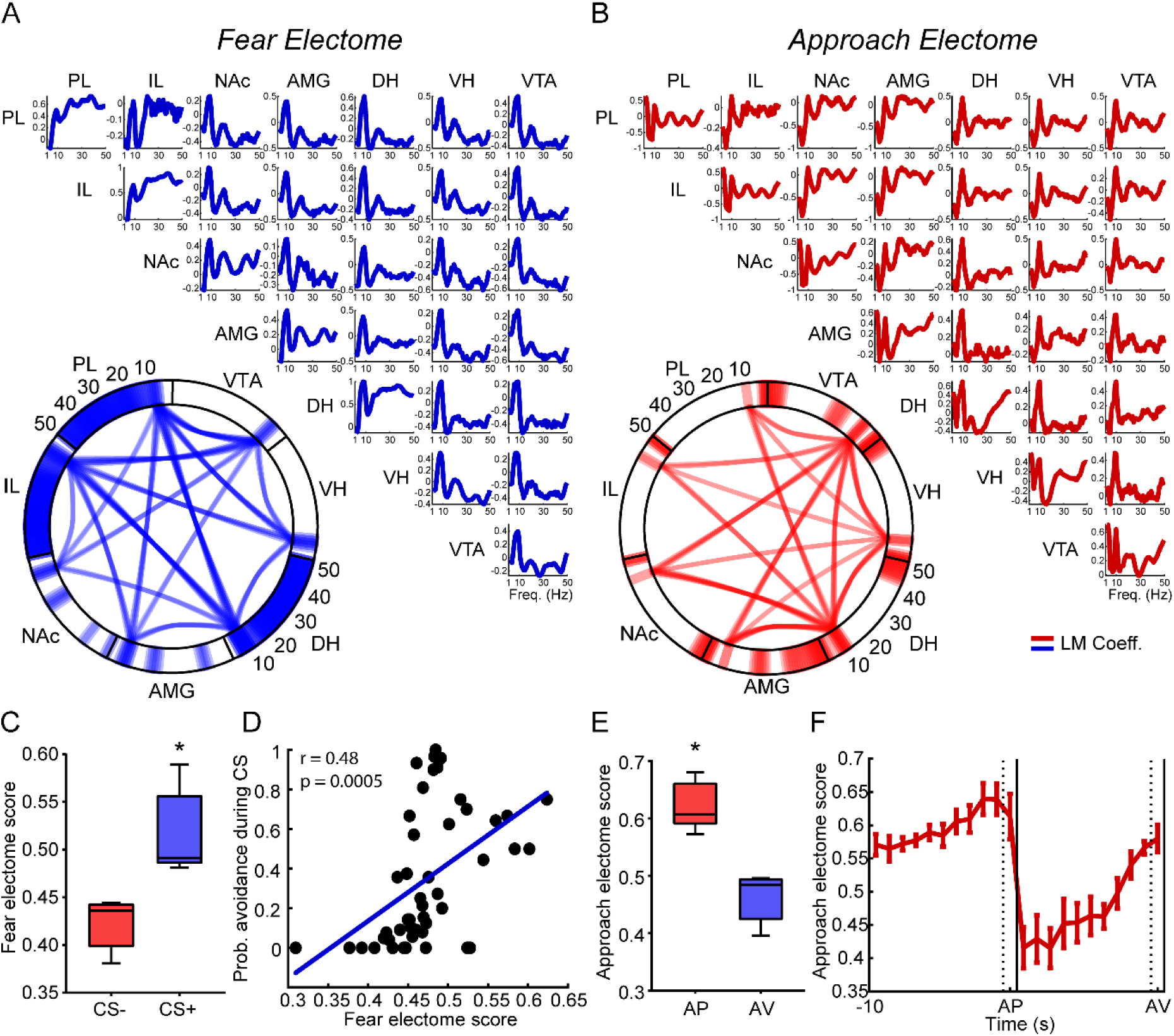
Patterns of brain-wide activity decode fear or safety and approach or avoidance across individuals. ***A-B*,** Electrical functional connectomes (electome) for decoding **(*A*)** fear vs. safety and **(*B*)** approach vs. avoidance. The models’ coefficients (lines) reveal a distinct positive relevance for brain-wide synchrony in the theta frequencies. ***C*,** Average fear electome scores discriminate CS- and CS+ in all rats. ***D*,** Fear electome scores correlate with probabilities of avoidance during CS on test sessions. ***E*,** Average approach electome scores discriminate the periods preceding approach (AP) and (AV) in all rats. ***F*,** Increases in the approach electome scores predict both AP and AV crossings (grid lines) but are especially strong preceding AP. *p < 0.05 (paired t-test).

In this regard, we realized that rats were usually on the reward during the most distant pre-AV epochs. Thus, we may interpret the low approach electome scores, not particularly as fear but also as a reward consummatory state. The same interpretation can be used for the robust power increase in the NAc in these epochs, which is a key region for reward processing. Again, the dynamics of decoder classification revealed behaviorally relevant information, which was only allowed at post decoding inspection. Therefore, the lower engagement of this pattern of brain-wide theta activity could be related to a consummatory state. In contrast, the increases of this activity could be linked to a motivational drive underlying both approach or avoidance, but particularly stronger in positive valence.

## Discussion

### An animal model to investigate fear, safety, approach, and avoidance

Here we implemented an experimental design by merging the protocols of learned safety and platform-mediated avoidance in a way that allowed the within-subject distinction of states of fear, safety, approach, and avoidance. The learned safety protocol is based on the work of Rescorla (1969, 1971), showing that while a CS that predicts aversive shocks eventually elicits fear responses, a stimulus that is explicitly unpaired with shocks suppresses them (summation) and is more difficult to associate later with fear (retardation). Our findings show that the CS-followed both criteria. It decreased the probabilities of avoidance, and CS+ took days to significantly increase them after reversal. Later studies in rodents showed that learned safety has antidepressant and stress mitigating properties, evokes distinct neural responses, and promotes approaches in anxiogenic environments, indicating these signals are not merely neutral (Pollak et al., 2008, 2010b). Our design differs from the original protocol in that we performed safety and fear conditioning in the same subjects instead of comparing two groups. It also differs from fear discrimination in that we conditioned an identical stimulus to both associations rather than two similar stimuli. In this regard, our work is the first investigation of oscillatory activity in this protocol of safety learning.

For the approach-avoidance task, we adapted the platform-mediated avoidance, designed by Quirk et al. (Bravo-Rivera et al., 2014; Diehl et al., 2019). Instead of using a Skinner box, we performed a low-cost open-source adaptation for reward delivery in a shuttle box. Although this protocol was originally designed to distinguish passive and active fear responses, we demonstrated that it could also reveal differential conditioning to fear and safety by increasing signaled avoidance and approach behaviors, respectively. In addition, this protocol clearly delineates positive (to reward) and aversive (to platform) valences of motivation to an identical behavioral response (crossing). We focused on active responses (avoidance and approach) rather than freezing to relate to the volitional aspect that defines stress coping (Lazarus and Folkman, 1984) and because interest in these processes has resurfaced in the last years (LeDoux et al., 2017).

### Decoding behavioral/cognitive states by oscillatory activity

Neural oscillations have been argued to provide a functional substrate while neuronal firing represents the actual code of information (Buzsáki et al., 2012). However, growing evidence shows that precise sensorimotor information can be decoded from electrical field potentials (Lebedev and Nicolelis, 2017; Anumanchipalli et al., 2019; Selfslagh et al., 2019; Frey et al., 2021). Here, we established a deterministic linear decoding pipeline based on standard machine learning methods and showed that multi-site LFP activities could decode stress coping-related states. The decoding performances depended on behavioral (CS association, learning curve, prediction of events) and biological (preference to theta frequencies and specific regions) variables. It exhibited significant generalization (across days in each rat) equivalently to ANNs, applicability for different investigations (fear vs. safety, and approach vs. avoidance), and allowed interpretation of the physiological relevance (a single coefficient to each variable). The pipeline used PCA for dimensionality reduction, which has been successfully used for neural decoding based on the firing of neuronal ensembles (Chapin et al., 1999; Narayanan and Laubach, 2009; Cunningham and Yu, 2014; Dejean et al., 2016) but also on LFP (Markowitz et al., 2011), including a recent study investigating subnetworks associated with depression in humans (Scangos et al., 2021).

### MLHFC theta network in stress coping

We found that a MLHFC network synchronizes in the theta frequencies, and the strength of this activity discriminates conditioned fear and predicts active responses to stressors, especially, approach. While we confirm previous findings of limbic-cortical theta synchrony during fear (Seidenbecher et al., 2003; Lesting et al., 2011; Likhtik et al., 2014), we also evidence the participation of the mesolimbic system in this “theta network”. Although theta oscillations are originated in the hippocampus (Buzsáki, 2002), all regions comprising this network have been reported to exhibit local neuronal firing phase-locked to theta field (Sirota et al., 2008; Royer et al., 2010; Adhikari et al., 2011; Van Der Meer and Redish, 2011; Kim et al., 2012; Likhtik et al., 2014).

We have recently shown that theta synchrony during fear is linked to the expectancy of control over stressors (Marques et al., 2022). Although we have previously investigated only the intermediate HPC, we found that the DH and VH also synchronize with the PFC in anticipation of escapable shocks. Although the VH is usually more associated with innate averseness, evidence indicates the DH is particularly engaged during learned fear (Çalışkan and Stork, 2019), agreeing with our findings. Our results also support that changes in corticolimbic directionality discriminate safety from fear (Lesting et al., 2013; Likhtik et al., 2014). In fact, directionality showed robust performance for decoding fear vs. safety, particularly within theta frequencies.

The link between theta and avoidance came from a series of studies using the elevated plus maze in mice, where increases in VH-PFC synchrony were linked to avoidance of the open arms (Adhikari et al., 2010; Padilla-Coreano et al., 2019). This task relies on rodents’ innate aversion to open areas, while the ‘approach’ motivation only occurs due to inherent exploratory drive. By making the reward component more explicit, we found that stronger theta activity predicted approach. Because theta activity is correlated with movement (McFarland et al., 1975), we do not discard the possibility that our findings on theta activity may be related to translocation, particularly during the conditioning sessions. However, in the AA task, we analyzed epochs that mostly considered preparatory states before crossings. Overall, these findings also fit a general proposition that theta correlates with preparatory rather than consummatory behaviors (Buzsáki, 2002).

In conclusion, our work suggests that theta activity underlies active stress coping, and this processing occurs at the brain-wide network level. The preferential engagement of network theta activity during controllable stress and positive motivational valence highlights the potential of theta modulating therapies for stress-related disorders characterized by cognitive and motivational deficits, such as depression. However, it suggests caution for those related to threat detection, such as anxiety and trauma.

### Brain-wide network processing

Large-scale brain networks have been implicated in cognition, emotion, and psychopathologies (Bressler and Menon, 2010; Menon, 2011; Bassett and Sporns, 2017). Identifying dysfunctional networks is a key goal to develop personalized treatments for anxiety and depression (Williams, 2016). Evidence that stress is associated with brain-wide activity comes from studies of resting-state neuroimaging (Magalhães et al., 2018, 2019), immediate-gene expression (Wheeler et al., 2013; Besnard et al., 2019; Worley et al., 2020), and, only recently, neural oscillations (Hultman et al., 2018; Scangos et al., 2021).

This is the first study of our knowledge to test the relationship between neural oscillations, the number of regions spread across a brain-wide network, and the encoding of stress-related information. We found a positive relationship between decoding performance of stress coping states and the number of regions included, which reached the best performance when all regions were considered. All regions performed better than each separately, even when considered as hubs. This relationship happened in a logarithmic function, which indicates that few regions already carry relevant information, but decoding still keeps improving with more regions. A similar pattern was found between decoding motor actions and the number of electrodes in the primate motor cortex, suggesting a generalized property of oscillatory coding (Markowitz et al., 2011). In addition, we also observed that all estimates – power, coherence, and directionality – presented greater performance when combined. Importantly, these findings cannot be explained by biases from a variable number of attributes because we fixed these numbers across comparisons. Thus, we suggest that the better performance of a brain-wide activity is evidence that stress coping-related information is encoded at a large-scale network level.

### Individual variability in brain synchrony and stress processing

Many reports show individual variability in oscillatory activity linked to stress processing (Likhtik et al., 2014; Hultman et al., 2018; Thériault et al., 2021; Marques et al., 2022). Here we found a significant generalization of decoder models tested within each rat but not between individuals, evidencing remarkable variability in stress processing. However, we observed that individual variability of oscillatory activity is greater for invariant brain activity between individuals than between stress states.

Synchrony was the measure with the most significant individual variability. We noticed whole-spectrum differences in coherence across pairs of regions. For example, most rats exhibited a markedly greater whole-spectrum coherence between the VTA and the NAc, while others did not. Adhikari et al. (2010) reported greater whole-spectrum coherence between the PFC and the VH than with the DH, suggesting this may be related to anatomical connectivity. If these variabilities result from the accumulated differences in the implant coordinates or if they emerge from brain differences remains elusive. Nonetheless, there is growing evidence that patterns of functional connectivity (Mueller et al., 2013; Finn et al., 2015) and electroencephalographic synchrony (Kondacs and Szabó, 1999; Kong et al., 2019) can be used to distinguish individuals, supporting a physiological basis. Also interestingly, despite the observed diversity of synchrony patterns, we noticed they all resembled a small-world topology, in agreement with previous studies on brain connectivity (Bassett and Bullmore, 2006; Bullmore and Sporns, 2012).

### Electomes

Machine learning is a powerful tool for identifying complex patterns in biological data, which often presents the drawbacks of difficult interpretation, reproducibility, and integration. Therefore, contemplating these aspects is required to ensure the trustworthiness of this approach (Vu et al., 2018; Heil et al., 2021). The electomes of fear and approach identified here are remarkably similar to the *electome factor 1* reported in Hultman et al. (2018). Both patterns are represented by increases in brain-wide theta synchrony and smaller increases in the beta ranges. Although power and directionality are distinct, the coherence similarity between patterns is close to qualifying this finding as a replication. Intriguingly, Drysdale et al. (2017) distinguished two functional connectivity patterns that correlated to anxiety or anhedonia measures, defining biotypes of clinical depression that are differentially responsive to treatments. Taken together, our findings indicate an important neurobiological distinction between the dimensions of motivation (valence/anhedonia) and threat detection (fear/anxiety vs. safety) that could guide neurophysiologically-informed treatment decisions for mood and anxiety disorders.

## Conclusions

Our work evidences that stress coping is processed at the brain-wide level with remarkable individual variability. Our findings also add to a recent proposition that theta oscillations underlie active stress coping, showing that it occurs at a large-scale network and is associated with both aversive and, especially, positive motivational valence. This study highlights the potential of multi-site electrophysiology combined with artificial intelligence to discover complex patterns of network function associated with stress-related disorders.

## Acknowledgements

This research was funded by Coordenação de Aperfeiçoamento de Pessoal de Nível Superior - CAPES - Finance Code 001 (D.B.M.: 88882.328283/2019-01; L.R.Z.: 88882.328297/2019-01), Conselho Nacional de Desenvolvimento Científico e Tecnológico - CNPq (J.P.L.: 305104/2020-9 and 422911/2021-6; T.P.: 164165/2018-5), and Fundação de Amparo à Pesquisa do Estado de São Paulo - FAPESP (D.B.M.: 2022/16812-3; R.N.R.: 2018/02303-4; M.T.R.: 2020/01510-6; B.A.M.: 2019/03445-0; J.P.L.: 2016/17882-4). We thank Luis Fernando Luca, Antonio Renato Meirelles e Silva, Renata Caldo Scandiuzzi, and Daniela Ribeiro for technical support. We also thank Adriano Bretanha Lopes Tort, Deborah Suchecki and Newton Sabino Canteras for discussions. DBM, RNR and MTR designed the research. DBM, MTR, BAM, TP, and LRZ developed the methodology. DBM conducted the experiments. DBM and RNR analyzed data. RNR and JPL supervised the project. All authors wrote, revised and approved the manuscript.

